# Evolution of the highest fidelity DNA replication systems

**DOI:** 10.64898/2026.04.16.719065

**Authors:** Stephan Baehr, Cooper Call

## Abstract

DNA mutation on average is deleterious, and evolution generally acts to reduce mutation rates to the limit of natural selection. The limit of natural selection is set by multiple factors, of which effective population size is only one. We consider lethal mutagenesis as an upper bound to mutation rates for any organism, an argument that is congruent with a biophysical context, wherein random mutations are a form of entropy. In this analysis, coding genome size, body mass, generation time, and temperature potentially explain more than 90% of the variation in mutation rate per generation across the Tree of Life. The organisms with larger genomes, longer lifespans and relatively larger body sizes, known and unknown, represent the lineages which have likely evolved novel mechanisms to lower mutation rates, a desirable trait for reducing cancer incidence. Though these variables are largely shared by Peto’s Paradox, this selective pressure responsible for the evolution of lower mutation rates occurs through germline mutation rate evolution rather than the soma.

**Significance:** DNA replication varies and is influenced by natural selection. Mutations are usually harmful and contribute to human diseases like cancer, Understanding the evolutionary factors behind DNA replication fidelity may help identify and develop novel strategies to reduce human mutation rates.

## Introduction

DNA mutation is the basis of all heritable biological variation. Some genetic variation gives rise to beneficial traits, most mutations are neutral or nearly neutral(1, 2), and some mutations result in severely deleterious phenotypes(3). Over evolutionary time, because average mutation is deleterious, selection generally acts to reduce organism mutation rates to the limit of natural selection, where one such limit, genetic drift, gives rise to the drift-barrier hypothesis (4–7). Effective population size is a constraint, but by no means the only constraint on the evolution of mutation rates. Specifically, although human mutation rates are among the highest measured across the tree of life(7), the human mutation rate per germline cell division is among the lowest(8, 9). Despite the census population sizes of humans being unarguably lower than that of bacteria, nevertheless, selection has acted to increase human DNA replication fidelity per heritable cell division. If humans are to seek novel methods of increasing DNA replication fidelity, where might we look and why?

Mutation rate evolution has been an outstanding curiosity to biologists for at least a century(6–8, 10–12). For example, there has been a long-recognized relationship between coding genome size and mutation rate(4, 6, 12); to avoid inheriting a deleterious mutation every single generation, larger genomes necessitate lower mutation rates. While the statement is accurate, a vexing problem of evolutionary biology is that many biological features co-vary; the organisms with larger genome sizes tend to also be larger and have longer lifespans/generation times and lower census population sizes. As generation times become longer, more spontaneous mutations arise(13–17), and often more cell divisions occur(8), increasing the mutation burden of the genome between selection events. Previous research has noted that natural selection’s ability to purge deleterious mutations breaks down when one or more deleterious mutations is inherited each generation(8, 18): if there is no significant chance of the “fittest” genotype replicating itself, Muller’s ratchet is invoked, a directional walk to mutational meltdown(19–25). To counteract the risk of mutational meltdown, recombination(8, 19, 20) allows a general means of escape from the ratchet, so long as underlying mutation rates are not excessive.

Evolutionary innovations that lower mutation rates evolve with some frequency(26, 27), though their best evidence of existence in natural populations comes from humans(28). In contrast to asexual species, recombining lineages suffer significantly less from the emergence of hypermutator mutations, but there is yet a fitness cost. For example, linkage of a small subset of the genome to a hypermutator allele will eventually incur some fitness cost which selection can “see”: if the fitness cost is higher than the organism’s effective population size. Because deleterious mutations frequently emerge with fitness effects on the order of 1% (1, 2, 29, 30), this analysis considers the perspective that significant individual linked mutations or summed mutation burdens to hypermutator alleles will influence mutation rate evolution of sexual species as well as asexual species. One expected outcome from this perspective is a reduction in mutation rates per germline cell division in larger, long-lived organisms. An exact theory explaining what magnitude of hypermutator or mutation burden evolution can see in recombining lineages, for what longevity or mutation burden, remains speculative, dependent ultimately on the fitness impact of mutation in addition to recombination frequency. At this time we simply hypothesize that certain biological factors generally influence how many mutations emerge between selective events: too many mutations emerging at some level becomes a problem, lethal mutagenesis. The lineages which resolve this problem persist over evolutionary time. We additionally note a certain congruence with evolutionary theory and biophysics with respect to mutation rate evolution. When life is interpreted as information and is contrasted by entropy, akin to the balance between selection and drift, the variables of physics extend to mutation rate evolution(31). Within a genome, information is stored in nucleotide sequence; the average coding mutation, besides incurring a fitness cost, removes information from a coding portion of genome, which was most often placed into it by natural selection(31, 32). By casting mutation rate evolution in terms of physics, the physical variables known to apply to entropy emerge in salience. The physics of entropy in DNA are established, and operate across biological scales. They are the similar variables(33): the number of random mutations accumulated (dependent upon lifespan, body size, and temperature), genome size, and the average information (or fitness) content per mutation. Temperature for example has been noted to affect DNA mutation rates(10, 34, 35). Notably, with respect to longer lifespan and larger body size, more mutations may accumulate, more information is potentially lost per generation in germline cells. Even with regard to somatic cells, post-replicative cells continue to accumulate DNA mutations, as do replicating cells, the latter at a seemingly somewhat faster rate.

Mutation rates are reported in units of base pair substitutions per site per generation, where generation is the unit of time, because that is the unit of time on which selection may act. For a human context of mutation–or entropy–as it may pertain to lifespan and cancer incidence, time in generations is unstandardized; a generation time may vary between 30 minutes(36) and at least 30 years(37, 38) across the Tree of Life. Therefore, we seek to understand mutation rate evolution in both contexts: mutation rate per cell division or year, and mutation rate per generation. It is well known that mutation rates vary across the Tree of Life; but the mutation rates which are the lowest per year or per cell division are perhaps more likely to provide molecular insights of how humans may lower their own mutation rates.

By examining mutation rate evolution, molecular biologists seeking novel mechanisms to lower human mutation rates may consider organisms of large coding genome size, generation time, and body mass. Regarding mutation rate per site *per generation*, this analysis finds that temperature along with body mass, generation time, and genome size are potentially sufficient to explain in excess of 90% of the variation in mutation rates measured.

## Results

### Mutation rate per standardized time

The question of which organism has the lowest mutation rate has often been asked, but the answer is often “depends”. Per generation, outliers include the single-celled eukaryotes *Paramecium*(39) and *Tetrahymena*(40), unicellular eukaryotes of unusual size. To remove the variable of generation time, we consider a perspective of mutation rate per year, converting generations into years. Figure 1 examines the relationship of organisms across the tree of life in mutations per year, in the context of coding genome size(Fig 1a) or body mass(Fig 1b). Notably, both variables exhibit a negative relationship across the tree of life. In Figure 1b, when only multicellular organisms are considered, the relationship between body mass and mutation rate per year approaches log linearity with an *r*^2^ value of 0.93. The lowest mutation rates per unit time remain the ciliates and the long-lived large organisms, *Paramecium tetraurelia* and the bow-head whale *Balaena mysticetus*.

**Fig. 1.**
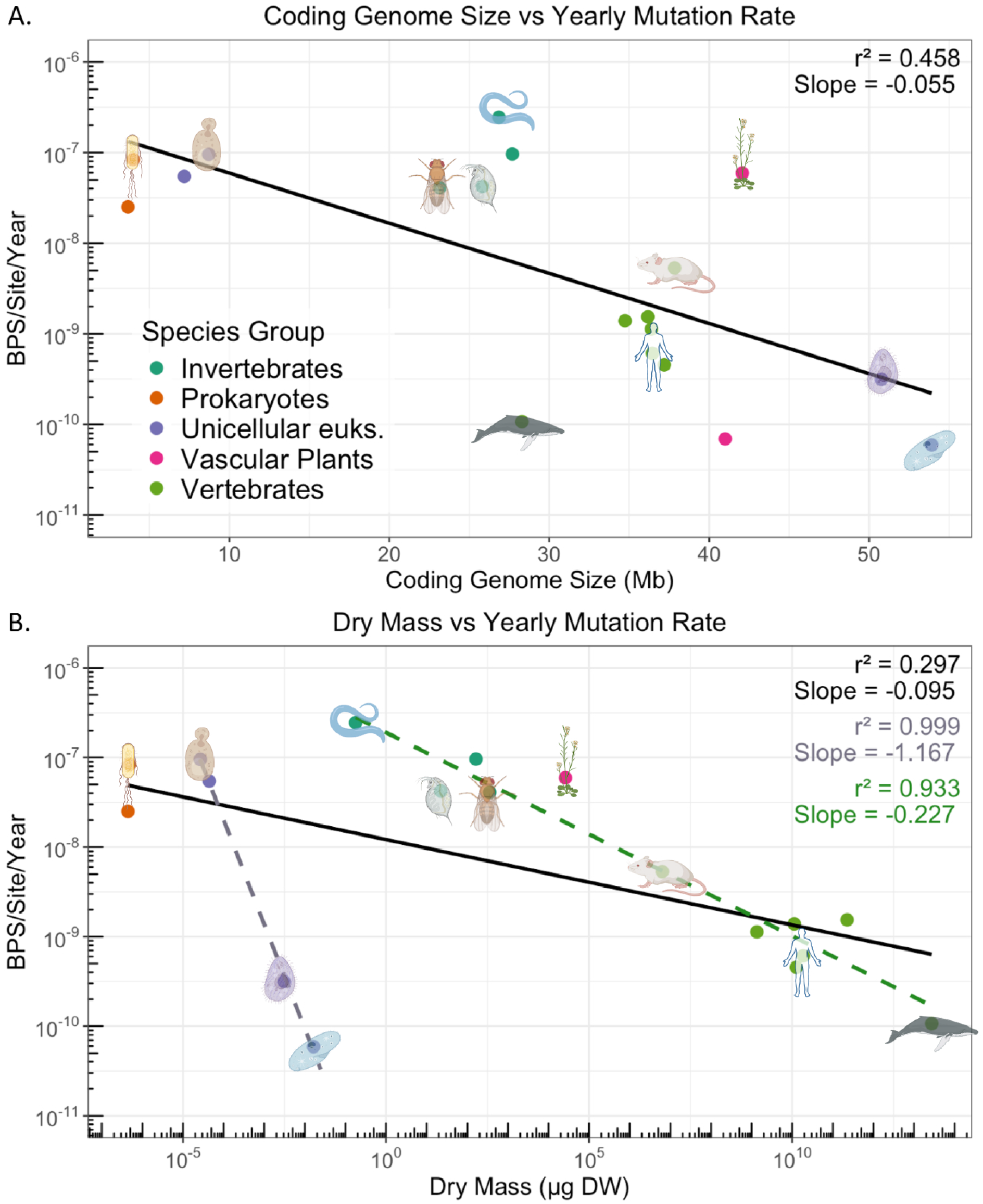
Mutation rate per year. Panel A: Mutation rates per year scale with coding genome size. Panel B: Mutation rates per year scale with dry mass across 20 orders of magnitude. Linear regressions for all organisms (black), multicellular eukaryotes (dotted green) and unicellular eukaryotes (dotted purple).

There is some criticism to the use of mutation rate per year, as mutations only arise somewhat as a function of time. For example, in the human germline the majority of mutations arise from the male lineage(17, 41–43), hypothesized to be due to recurrent cell divisions(8). Therefore, it is also possible to convert mutation rates per generation into mutation rates per germline cell division. For unicellular eukaryotes, these numbers are the same. Multicellular eukaryotes require some knowledge of anatomy and developmental patterning, which is typically present for only the most well-established model organisms. Coding genome size appears a strong predictor from this limited dataset, Figure 2a. Surprisingly, over 20 orders of magnitude of body mass yield only a two-order variation in mutation rate per germline cell division, and dry mass is only weakly predictive of germline mutation rates, with an *r*^2^ of 0.09, Figure 2b. We note that in the 28 years since an analysis of mutation rate per germline cell division(8), only one additional multicellular organism’s germline cell division count has been estimated, an oversight by the field(44). Regardless, by this analysis, the ciliates remain the organisms with the lowest mutation rates per germline cell division. By the analyses of either Figure 1 or Figure 2, considering time in terms of years or germline cell divisions, bacteria and yeast have among the higher mutation rates across the tree of life. Selection may not act upon the timescales described here, but the trends seen remain relevant to questions of where to look when seeking novel mechanisms to reduce human disease incidence.

**Fig. 2.**
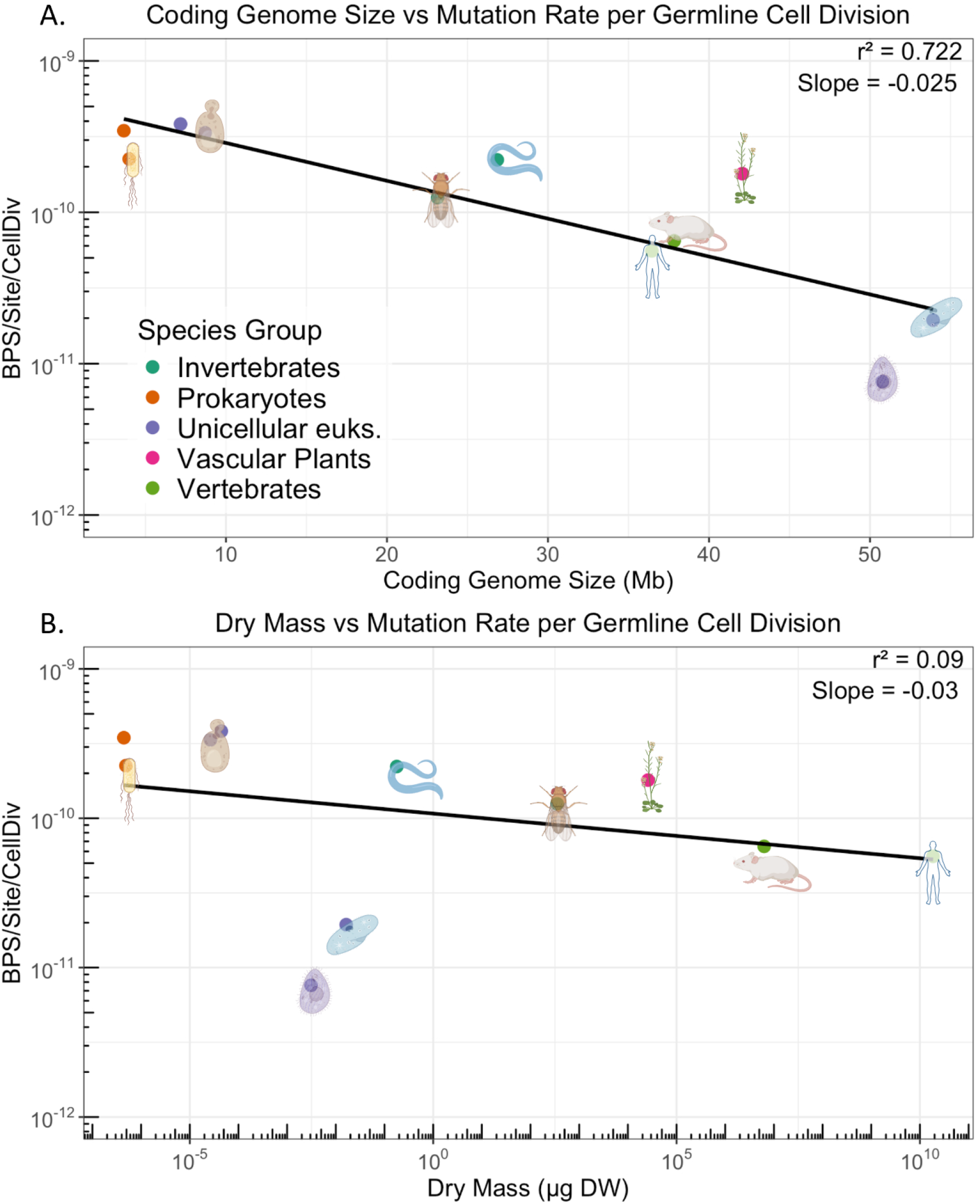
Mutation rate per cell division. Panel A: Coding genome size vs. mutation rate per germline cell division. Panel B: Dry mass vs. mutation rate per germline cell division.

### Mutation Rate per Generation

The evolution of mutation rates per generation is correlated with various features. Previous analyses within *mammals* has suggested a majority of the difference mutation rates between organisms is due to generation time alone(45); but across the tree of life as shown within this article’s dataset, the *r*^2^ value remains particularly weak at 0.04 (Figure 3). Dry mass and coding genome size exhibit similar performance issues, though coding genome size perhaps the strongest at a *r*^2^ of 0.2. Rather than attempting to inflect these data by invoking effective population size as previous research has considered(6, 7), we additionally consider the context of increasing mutational burden per generation and its similarity to entropy.

**Fig. 3.**
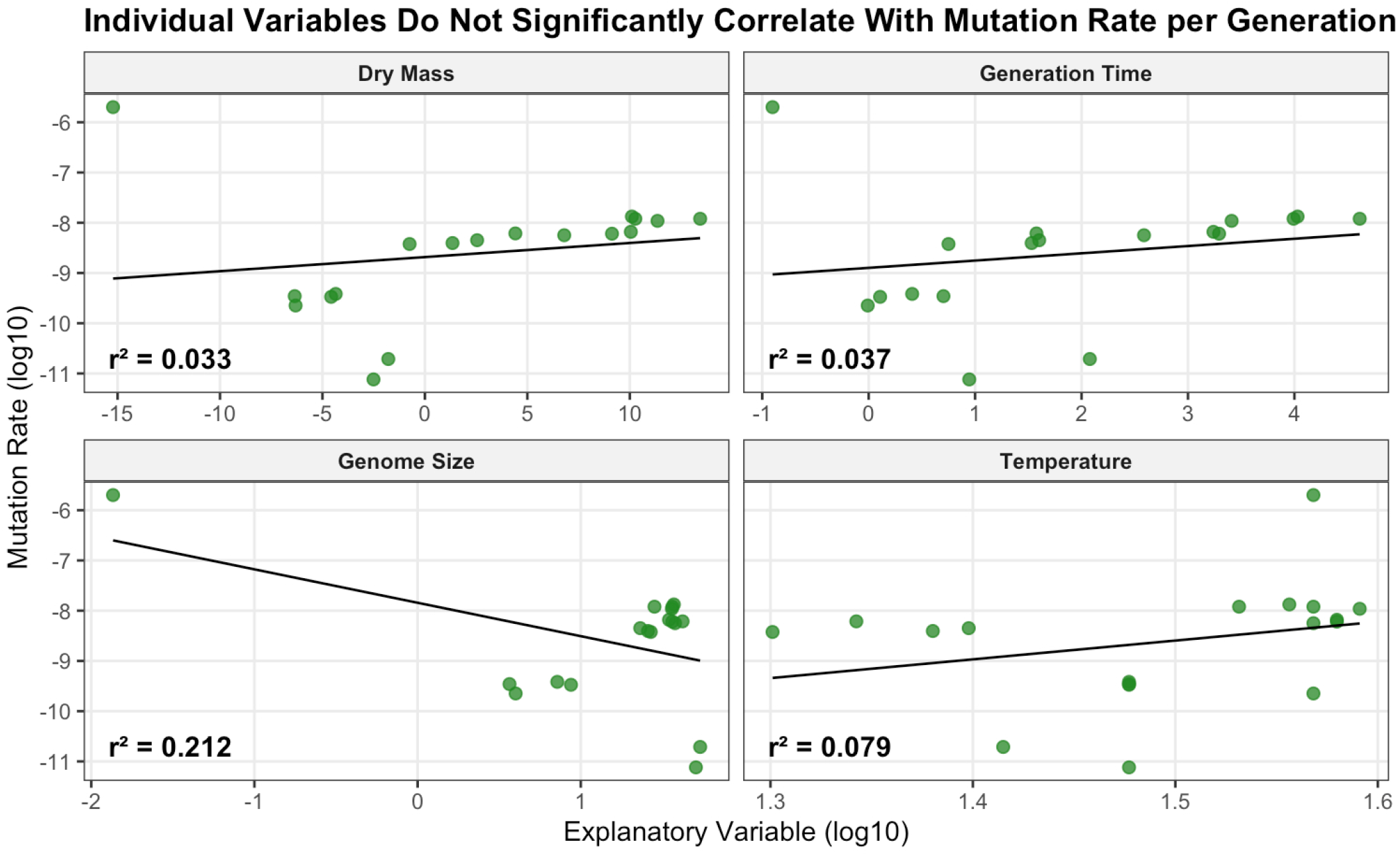
A biophysical perspective of mutation rate evolution. Physical parameters of organisms in this dataset alogn correlated poorly with base pair substitution rate per site per generation. Genome size refers to coding genome size.

The factors which describe the number of mutations which arise per generation are body size, lifespan, temperature, and the total hit-size of the genome. Absent knowledge about how the fitness impact of mutations may vary between organisms, these are the best variables available that describe information theory entropy and its accumulation over time in DNA. The number of mutations accumulated in a germline cell in a generation is influenced by lifespan, temperature, the number of bases to be replicated, and how many cell divisions must occur between selection events. These four variables together in a 5-dimensional multiple regression upon mutation rate per generation (Figure 4) yield a raw *r*^2^ value of 0.926. While initially satisfying, several statistical impediments hinder acceptance of this number at face value. The variables listed above notoriously covary, and the dataset of 19 organisms is insufficient to entirely gauge the validity of the model; the rule of thumb of 10 observations per explanatory variable is not met. The number of observations is difficult to meaningfully increase; these 5 descriptive statistics are challenging to obtain across the Tree of Life. In an example, though there are many bacterial mutation rate estimates, only two(36, 46) estimates of bacterial senescence have as yet been rigorously quantified. To account for overfitting and co-variance, we examine the model with a Leave One Out Cross-Validation. An organism was removed from the model, and the model was tested to observe how accurately it could predict the missing organism’s mutation rate. This was reiterated for all of the organisms to obtain a cross-validation value. The cross-validation *r*^2^ is 0.745. This demonstrates that some overfitting is present in the model, but even when taken into account, the model had a significant amount of predictive ability for mutation rates, *p* < 2.7 × 10^−8^.

**Fig. 4.**
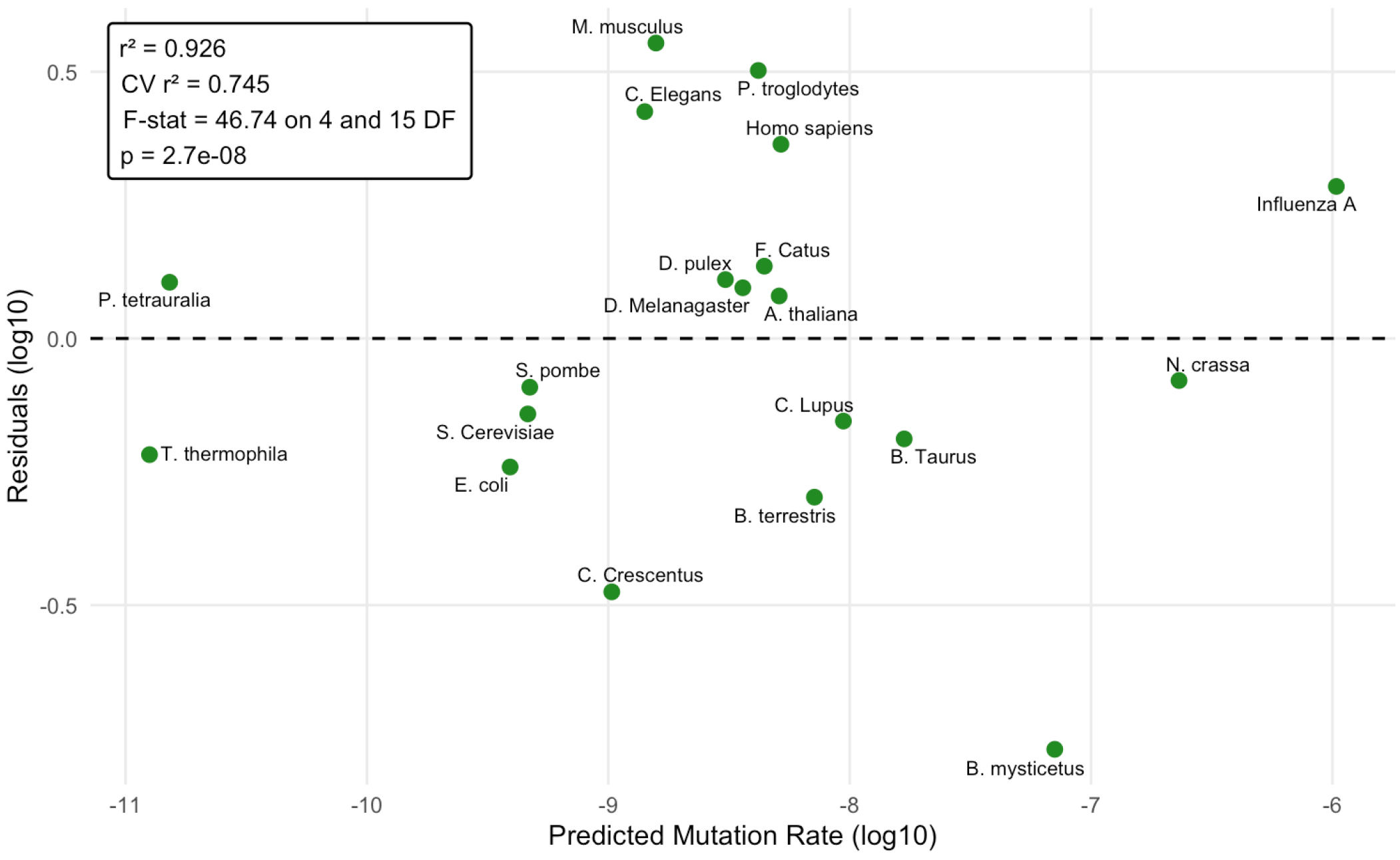
Aggregate parameters are predictive of mutation rates per generation. A residual value of ±1 represents a deviation of an order of magnitude from the models predicted mutation rate. A residual value of ±0.5 represents approximately a three fold deviation. The residual plot reveals that all but two points are below this threshold and all organisms were within an order of magnitude of the linear model.

In Figure 4, the variance of mutation rates across the Tree of Life per generation are plotted against the residuals of the multiple regression in a log-log plot. For this graph, perfect correlation between the features is present upon a dotted central line. A residual plot reveals the average magnitude of each organisms deviation from the predicted linear model. Upon such widely variant mutation rates, genome sizes, and body masses, scaling 5, 6 and 22 orders of magnitude, the variance between the per-generation mutation rates measured and the model’s estimate vary by 1 order of magnitude or less. Absent the outlier of *B. mystecetus*, notably the least-rigorous datapoint from a single whale trio(47), most lineages fall satisfyingly closer to the mark, resulting in an *r*^2^ value of 0.96 and a cross-validated *r*^2^ of 0.86. We retain the bowhead whale data point due to its placement alongside other whales providing some additional confidence, and also for its potential importance to understanding Peto’s paradox. The formula for the fitted linear model is as follows (Equation 1):

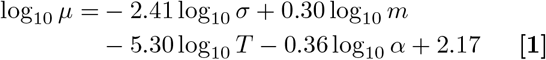

Where *µ* is mutation rate (mutations/site/generation), *σ* is effective genome size (Mb), m is dry mass (*µg*), T is temperature (*C*), and *α* is generation time (days). The F-Stat was 40.76 with an associated p-value of 3.09 × 10^−7^. To give an example of a calculation for the model, consider humans. Humans have a coding genome size of 36.45 Mb, an average dry mass of 1.86 × 10^10^*µg* (7), an average internal body temperature of 37^*°*^C, and a generation time of 9818.5 days(37). The model predicts a mutation rate of 5.7 × 10^−9^. Our expected value, the known human mutation rate, is 1.2 × 10^−8^(9), a 2.1-fold difference.

To aid the understanding the multidimensional nature of the data, the dimensions have been transposed into a more intuitive graph in Figure 5. Per-generation mutation rate, the dependent variable, was plotted on the z axis. The x and y axes are assigned with effective genome size and generation length. The size of the individual points represents the dry mass of the organism, on a log-scale, and the color represents the temperature for which the organisms mutation rate was measured. This multidimensional object demonstrates how discrepancies in certain dimensions can be accounted for by others; how weak individual variables can in sum account for the mutation rates empirically measured.

**Fig. 5.**
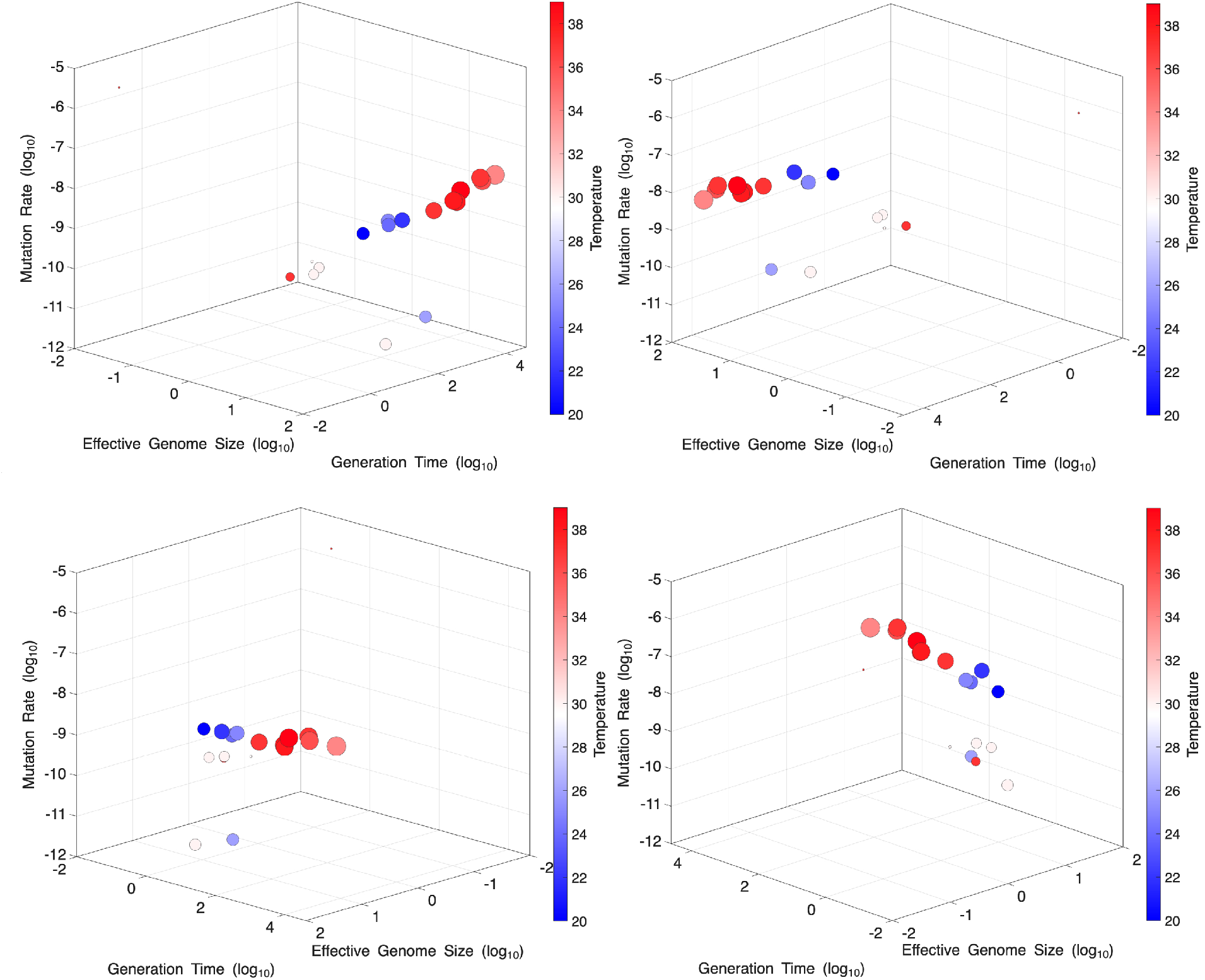
Five-dimensional data visualization of mutation rate evolution. Multi-dimensional perspectives allow data points that appear as outliers in a single dimension to be contextualized through additional variables. This object, viewed in four perspectives, shows how incongruous data points like the ciliates (recurrent two outliers) are not explained from some perspectives but do align at others. Specifically, although the ciliates are much smaller than humans, they appear to undergo a similar number of germline genome replications between meiotic events with relatively large coding genomes.

## Discussion

This manuscript examines the parameters which explain variance in genetic mutation rates per unit time where time is considered per year, per cell division, and per generation. Evolution acts on the latter, but the first two have some relevance to impacting phenotypes over human lifespans. In this analysis, a certain congruence of evolutionary biology and biophysics provides a rationale for the parameters considered. Random mutation absent selection on average incurs a fitness cost(2); random mutation absent selection is also an expression of entropy(31). The parameters of body mass, generation time, genome size, and organism temperature influence the rate of propagation of genetic mutation, holding generally across all known biological scales of life, from viruses to whales. When examining mutation rates across the Tree of Life, it is clear that certain parameter spaces are biophysically possible but are not occupied, a survivorship bias. For example, if humans had the same per-cell mutation rate as *E. coli*, Figure 2a, instead of gaining ∼100 mutations per generation humans would have ∼300 mutations per generation, or at least 3 coding mutations per generation on average. We can infer that the reason such human lineages are not seen on average is that they are evolutionarily inviable(18). By this perspective, the selective cost to excess mutation is visible to even the human or whale lineages, despite limited effective population sizes relative to mice, fruit flies, or bacteria. Hypermutator strains must continually arise and be continually be eliminated by purifying selection over evolutionary timeframes. On the other side, mutation rates are not lower because of the expense of DNA repair when unneeded is prohibitive(18) and its benefit is beyond selection’s resolution(48).

With regard to effective population size *N*_*e*_, some selective resolution is required to forestall mutational meltdown. Proponents of effective population size quickly note that the variables chosen for examination here, such as dry mass and generation time, also strongly co-vary with effective population size(49); is it not simply that effective population size is conferring the true signal which determines mutation rates? Without disputing the importance of effective population size in conferring a drift barrier, the strict dependence upon knowing mutation rates to estimate *N*_*e*_ creates several problems. Admittedly, the paradigm of *N*_*e*_ perhaps appears satisfying at first glance. However, *N*_*e*_’s strong dependence upon mutation rate would estimate that there are on the same order of magnitude of effective bacteriophages, RNA viral species with small genomes, as there are effective humans; this is a challenging conclusion to accept, given that viral particles are estimated to outnumber bacterial cells by at least an order of magnitude. Further, the equation used for calculating *N*_*e*_, *θ* = 4*N*_*e*_*µ*, holds among its assumptions that a population be at equilibrium. Older species appear to have more diversity than younger species(50, 51), which predispose different estimates of *N*_*e*_. Sites for *N*_*e*_ estimates are also assumed to be evolving neutrally, despite evidence of selection at linked sites for much of the genome(52, 53). Sidestepping these concerns is a physical or lethal mutagenesis understanding of mutational hazard and the range of viable solutions for organisms of varying genome size, generation time, temperature, and a proxy of germline cell division count via dry mass. A significant strength of this study is that the model appears sufficiently flexible to encompass all organisms that replicate their genomes, including viruses. This model is a falsifiable, testable hypothesis that bears further testing; as an aside, being *falsifiable* is something that measures of *N*_*e*_ inherently are not. Regardless, the model offered here is congruent to *N*_*e*_ overall, rather than in direct competition. Greater selective resolution is perhaps helpful, but by no means required to build the highest fidelity DNA replication systems.

In the pursuit of outlier organisms exhibiting low DNA mutation rates, researchers have considered Peto’s paradox(54). Given that germline mutation rates per cell division are significantly lower than somatic mutations, it seems unlikely that in most lineages that somatic cancer burden is a primary driver of DNA mutation rate innovation. In other words, Peto’s paradox is somewhat facile in its original interpretation, ignoring the significantly greater problem of per-generation germline mutation rates relative to the emergence of cancer at old age, when selection is relatively weak(55). By an evolutionary perspective, cancer is a minor problem, something to keep in mind in a human-centric, and particularly an aging human world. Recent interpretations of cancer prevalence in mammals suggests that larger organisms on average do tend to incur more cancer incidence, abrogating the paradox(56). As researchers note, the same DNA replication systems(6) that influence germline mutation rates are those that influence somatic mutation rates, a genetic linkage which may prove sufficient to explain the variance in somatic mutation rates decreasing in larger, long-lived mammals(45).

A fundamental issue to the pursuit of lowering human mutation rates regards the somatic and germline divide. Evolutionary theory on the disposable soma(55, 57) adequately explains why the bifurcation exists; why somatic mutation rates per cell division are so much higher. Less clear is how researchers might take the strategies of germline cells back to the soma. For example, research indicates that *Paramecium*(58) also adhere to the disposable soma, featuring macronuclear mutation rates higher than their micronuclear mutation rates; but if so, even their somatic mutation rates per cell division appear to be lower than humans by several-fold per cell division. This line of inquiry does lead to the question of what a generation is in Ciliates with macronuclei: if a generation is the time between germline mutations being exposed to selection, the per-generation mutation rate of *Paramecium* and *Tetrahymena* may need to be adjusted. In any case, questions as to whether the germline strategies are exactly tangible to the human soma, the germline rates provide a hint as to which organisms likely have lower somatic mutation rates per unit time(58).

Under the perspective given here, it may prove challenging to lower germline human DNA mutation rates, whether by behavior, drug, or by genetic modification. Contrary to the perspective that human mutation rates are high and there are many changes to be incorporated to improve our DNA replication systems, spanning orders of magnitude, humans display relatively low DNA mutation rates under any perspective of time other than per-generation. As for somatic mutation rates, it may be possible to simply look to our own genomes and germline lineages in addition to venturing to other organisms. Some initial work on outlier organisms has already begun, to considerable interest and promise(59–62). The increase in average human generation time, which already has occurred and may continue to occur, may eventually make such interventions attractive or else risk an increase in human genome mutational load(24, 25).

There are several limitations to this study. There are many more mutation rates that have been measured(7) than there are concrete estimates of germline cell divisions or even lifespans, as reported here. Others have measured mutation rates per unit time(58, 63, 64) but the methodologies do vary somewhat. Of particular note, the replicative senescence of bacteria is challenging to score, needing strain-specific quirks or semi-bespoke “mother machine” microfluidic devices(36). A greater study of bacteria and their replicative longevity would allow for greater analysis within and between diverse clades of life. It is not clear if or why bacteria may be influenced differently from multicellular eukaryotes by the variables of dry mass or effective genome size. The same may also be said for viruses, unicellular eukaryotes, or plants, which do not feature prominently in this study. With regard to unicellular eukaryotes, there are only four accounted here; dry mass does scale with mutation rate per year, with an *r*^2^ of 0.99 Figure 1b. Given the paucity of data, at this time we can’t put much weight to the regression. To get around limited composite sample size of organisms with known values for all variables, other groups have turned to proxies or regressions of one variable which is measureable(CITE EYRE WALKER), for example body size vs. what is not known or readily measurable, such as replicative lifespan of a bacterial pole. To avoid over-reliance on assumptions, we report the trend only from the real data we can reference.

With respect to linear regression in evolutionary biology, a common concern regards phylogeny itself masking or otherwise modifying the interpretation of data(56, 65, 66). We considered if phylogeny is obscuring the trend of the data under analysis of this manuscript. It is true that humans relatively oversample the clades of life which are visible to humans: large and especially mammalian.

However, at this time the overall range of the data, from viruses to Bowhead whales, appears robust to the possible interference of phylogeny.

One incidence worth some comment is how remarkably *good* the estimates of mutation rates have historically been, on average, even as technology has dramatically advanced. The estimates summarized by Drake et al., 1998 (8) for example have held up remarkably well, moreso perhaps than the authors may have envisioned at the time of the document’s writing. Even the initial estimates of temperature’s influence on mutation rate by Muller reported in 1928(10) have held up remarkably well in relation to modern experimentation(34, 35), as have the human mutation rate estimates of Haldane in 1935(67). These earlier investigators obtained experimental estimates of mutation rates, and with conscientious analysis, they arrived at figures quite near those of contemporary mutation accumulation and trio-type experiments with single base-pair resolution.

We hope that this study provides further credence to the study of long-lived, large, and otherwise unique organisms as to how the lineages have found solutions to their mutational environments. The paper provides clear criteria in the identification of potentially exceptional organisms, even those as yet unknown or uncharacterized: coding genome size, for example. It may be that there are other ciliates or unicellular eukaryotes, as-yet unidentified, with large coding genomes. Also, this study highlights large gaps in aging research, for example in the realm of long-lived plants, in the ascertainment of their mutation rates and other molecular-biological features.

## Methods

### Estimation of generation time in years

A significant challenge in the search for low mutation rates per year is the estimation of what a generation is in standard time units. A generation time may vary, depending upon the organism, by over an order of magnitude. In humans, a generation time can vary from 14 years to 50 years in age; but mice generation time of 2 months to 2 years. Selection for the most part can only act upon organisms capable of reproduction, and therefore, consider the average of reproductive lifespan among females in sexual species as a proxy for a measurement of lifespan. This proxy is chosen in large part because statistics have been measured in many organisms, from model organisms to farm animals to humans, which allow its calculation.

The minimum lifespan is the average age of reproductive maturity, and the maximum lifespan is the point where half of females are no longer capable of reproducing. Among sexual species, males tend to have reproductive capabilities for a longer time than females, as is true in humans; a weighted average would skew average lifespans longer if male fertility were taken into account. However, decreasing fertility rates(68) with age and increased mortality rates among older mothers/fathers is a variable which would reduce average lifespan over evolutionary time relative to modern biological limits. By choosing the limiting reproductive lifespan among sexual species, there is an overestimation of older mothers contributing to lifespan and an underestimation of older fathers contributing to lifespan which for the most part negate each other. The test case of humans is a fair example: the average generation time of humans over evolutionary time has been approximated from mutation spectra data, to about 26.9 years(37), an average that is higher than many researchers initially expect. The high average arises because though reproductive viability begins around 13 years for females and child/maternal mortality are high, many human parents were capable of generating multiple offspring over a lifespan. Additionally, mismatches in paternal age and maternal age were common in pre-modern societies: human males could frequently range between 2 to 50 years older than their female partners. Though the “age gap” of such extremes is now generally culturally unacceptable, it remains a part of humanity’s evolutionary past. Using the contrived proxy of an average of average sexual maturity and average age of menopause, 13 and 50 respectively, The average reproductive becomes 31.5 years, fairly close to an empirically calculated 26.9 years; better than choosing the lower or upper value alone. For order-of-magnitude comparisons the accuracy of the calculation is sufficient. This calculation of average reproductive age is more accurate than simply choosing the first or last reproductive events as a proxy for lifespan, or the outlier events of oldest living member of any organism. Surprisingly, the linear regressions did not significantly change when calculating lifespan as the oldest recorded organism of a species, for example 123 years instead of 26.9 years for humans. This insignificance is due to lifespans of the dataset spanning four orders of magnitude.

Among non-sexual species such as *E. coli*, the average lifespan is calculated as the average between the first reproductive event and the time-point where half of individuals are no longer able to reproduce, or the average time to replicative senescence. These measurements have only been made for a few model organisms, and we hope that in the coming years more measurements of replicative senescence will be forthcoming.

### Estimation of mutation rate per mitosis

The estimation of the number of cell divisions per germline per generation is a challenging task that has languished, despite improvements in certain methodologies, to track cell divisions in development. Developmental biologists identified estimates of germline cell divisions 30 years ago, and additional estimates remain elusive. We have applied modern estimates of mutation rates to known estimates of germline cell division counts.

### Multivariate linear regression

A multiple linear regression was preformed on the data with mutation rate as the dependent variable and effective genome size, dry mass, temperature, and generation time as the explanatory variables. The linear model was fit using the lm() package in RStudio. The adjusted *r*^2^, F-test, and p-value were calculated using the summary() package. The F-test was preformed on 4 and 13 degrees of freedom; *df*_*reg*_ was 4: 4 explanatory variables, and *df*_*res*_ was 13: 17 species minus 4 explanatory variables minus one. The p-value was calculated using an F-distribution with the summary() package. Prior to running this correlation, we decided to test temperature, generation time, dry mass, and effective genome size and only these variables. A variety of variables with a possible connection to mutation rates has been discussed by the evolutionary biology community(69); we selected these particular variables for their influence and parallels in thermodynamics and entropy. We did not have to correct beyond the cross-validation *r*^2^ for multiple variables since only a single set of variables was considered. Since no variables were added or discarded to the linear model, further adjustment for multiple testing was not necessary.

## Supporting information

Supplementary References

## Acknowledgments

The authors wish to thank their advisor Michael Lynch for recurrent comments and review, and for providing a stimulating educational environment. We also thank Citlalli Alvarez for her efforts to begin this conversation. This work has been funded in part by NIH GMS grant 5R35GM122566-08 and the National Science Foundation, DBI-2119963, 2021-2026, BII: Mechanisms of Cellular Evolution. Figures 1 and 2 were created with the assistance of Biorender.

## Data, Materials, and Software Availability

The dataframe used to derive this analysis can be found in the supplemental information(.xls). The primary resource for mutation rate data arose from(7). The scripts associated with the construction of the graphs can be found at https://github.com/Coopergabriel-research/2026-Evolution-of-the-Highest-Fidelity-Replication-Systems

## SI Datasets

The Supplementary Information containing the references and parameters of the plots can be found in the attached PDF.

An excel file is also attached, containing the parameters by a more useable format.

