## Supplementary References for "Evolution of the highest fidelity DNA replication systems"

April 11, 2026

Table 1: Biological Parameters Across Diverse Species

| Species | Measurement Type | Reference |
|---|---|---|
| **D. pulex** | mutation rate | Lynch et al. (2023) |
| | age first reproduction | Lynch (1984) |
| | age last reproduction | Lynch (1984) |
| | generation time | N/A |
| | temperature | Keith et al. (2016) |
| | # of cell divisions | N/A |
| | dry mass | Lynch et al. (2023) |
| | genome size | Lynch et al. (2023) |
| **D. magna** | mutation rate | Lynch et al. (2023) |
| | age first reproduction | Lynch (1984) |
| | age last reproduction | Lynch (1984) |
| | generation time | N/A |
| | temperature | N/A |
| | # of cell divisions | N/A |
| | dry mass | Lynch et al. (2023) |
| | genome size | Lynch et al. (2023) |
| **S. cerevisiae** | mutation rate | Lynch et al. (2023) |
| | age first reproduction | Sherman (2002) |
| | age last reproduction | Minois et al. (2005) |
| | generation time | N/A |
| | temperature | Parapouli et al. (2020) |
| | # of cell divisions | N/A |
| | dry mass | Lynch et al. (2023) |
| | genome size | Lynch et al. (2023) |
| **H. sapiens** | mutation rate | Lynch et al. (2023) |
| | age first reproduction | N/A |
| | age last reproduction | N/A |
| | generation time | Wang et al. (2023) |
| | temperature | Geneva et al. (2019) |
| | # of cell divisions | Drost and Lee (1995) |
| | dry mass | Lynch et al. (2023) |
| | genome size | Lynch et al. (2023) |
| **M. musculus** | mutation rate | Lynch et al. (2023) |
| | age first reproduction | De Magalhães and Costa (2009) |

**Table 1 – continued from previous page**

| Species | Measurement Type | Reference |
|---|---|---|
| | age last reproduction | De Magalhães and Costa (2009) |
| | generation time | N/A |
| | temperature | Mei et al. (2018) |
| | # of cell divisions | Drost and Lee (1995) |
| | dry mass | Lynch et al. (2023) |
| | genome size | Lynch et al. (2023) |
| **C. elegans** | mutation rate | Lynch et al. (2023) |
| | age first reproduction | Zhang et al. (2020) |
| | age last reproduction | Zhang et al. (2020) |
| | generation time | N/A |
| | temperature | Flavel et al. (2018) |
| | # of cell divisions | Denver et al. (2009) |
| | dry mass | Lynch et al. (2023) |
| | genome size | Lynch et al. (2023) |
| **C. lupus** | mutation rate | Lynch et al. (2023) |
| | age first reproduction | N/A |
| | age last reproduction | N/A |
| | generation time | Mech et al. (2016) |
| | temperature | Kreeger et al. (1990) |
| | # of cell divisions | N/A |
| | dry mass | Lynch et al. (2023) |
| | genome size | Lynch et al. (2023) |
| **F. catus** | mutation rate | Lynch et al. (2023) |
| | age first reproduction | De Magalhães and Costa (2009) |
| | age last reproduction | De Magalhães and Costa (2009) |
| | generation time | N/A |
| | temperature | Giannetto et al. (2022) |
| | # of cell divisions | N/A |
| | dry mass | Lynch et al. (2023) |
| | genome size | Lynch et al. (2023) |
| **E. coli** | mutation rate | Lynch et al. (2023) |
| | age first reproduction | Gibson et al. (2018) |
| | age last reproduction | Wang et al. (2010) |
| | generation time | N/A |
| | temperature | Lee et al. (2012) |
| | # of cell divisions | N/A |
| | dry mass | Lynch et al. (2023) |
| | genome size | Lynch et al. (2023) |
| **C. crescentus** | mutation rate | Lynch et al. (2023) |
| | age first reproduction | Wright et al. (2015) |
| | age last reproduction | Ackermann et al. (2003) |
| | generation time | N/A |
| | temperature | Long et al. (2018) |
| | # of cell divisions | N/A |
| | dry mass | Lynch et al. (2023) |
| | genome size | Lynch et al. (2023) |

**Table 1 – continued from previous page**

| Species | Measurement Type | Reference |
|---|---|---|
| **P. tetraurelia** | mutation rate | Lynch et al. (2023) |
| | age first reproduction | Sung et al. (2012) |
| | age last reproduction | Takagi et al. (1987) |
| | generation time | N/A |
| | temperature | Luhring and DeLong (2016) |
| | # of cell divisions | N/A |
| | dry mass | Lynch et al. (2023) |
| | genome size | Lynch et al. (2023) |
| **T. thermophila** | mutation rate | Lynch et al. (2023) |
| | age first reproduction | Seyfert et al. (1984) |
| | age last reproduction | Brito et al. (2010) |
| | generation time | N/A |
| | temperature | Long et al. (2013) |
| | # of cell divisions | N/A |
| | dry mass | Lynch et al. (2023) |
| | genome size | Lynch et al. (2023) |
| **P. troglodytes** | mutation rate | Lynch et al. (2023) |
| | age first reproduction | De Magalhães and Costa (2009) |
| | age last reproduction | De Magalhães and Costa (2009) |
| | generation time | N/A |
| | temperature | Jensen et al. (2009) |
| | # of cell divisions | N/A |
| | dry mass | Lynch et al. (2023) |
| | genome size | Lynch et al. (2023) |
| **D. melanogaster** | mutation rate | Lynch et al. (2023) |
| | age first reproduction | Fernández-Moreno et al. (2007) |
| | age last reproduction | Piper and Partridge (2016) |
| | generation time | N/A |
| | temperature | Mołoń et al. (2020) |
| | # of cell divisions | Drost and Lee (1995) |
| | dry mass | Lynch et al. (2023) |
| | genome size | Lynch et al. (2023) |
| **A. thaliana** | mutation rate | Lynch et al. (2023) |
| | age first reproduction | Boyes et al. (2001) |
| | age last reproduction | Boyes et al. (2001) |
| | generation time | N/A |
| | temperature | Rivero et al. (2014) |
| | # of cell divisions | N/A |
| | dry mass | Lynch et al. (2023) |
| | genome size | Lynch et al. (2023) |
| **B. taurus** | mutation rate | Lynch et al. (2023) |
| | age first reproduction | De Magalhães and Costa (2009) |
| | age last reproduction | De Magalhães and Costa (2009) |

**Table 1 – continued from previous page**

| Species | Measurement Type | Reference |
|---|---|---|
| | generation time | N.A |
| | temperature | Sammes et al. (2019) |
| | # of cell divisions | N/A |
| | dry mass | Lynch et al. (2023) |
| | genome size | Lynch et al. (2023) |
| **P. sitchensis** | mutation rate | Lynch et al. (2023) |
| | age first reproduction | Trojan et al. (2026) |
| | age last reproduction | Trojan et al. (2026) |
| | generation time | N/A |
| | temperature | N/A |
| | # of cell divisions | N/A |
| | dry mass | Lynch et al. (2023) |
| | genome size | Lynch et al. (2023) |
| **S. pombe** | mutation rate | Lynch et al. (2023) |
| | age first reproduction | Vyas et al. (2021) |
| | age last reproduction | Spivey et al. (2017) |
| | generation time | N/A |
| | temperature | Farlow et al. (2015) |
| | # of cell divisions | N/A |
| | dry mass | Lynch et al. (2023) |
| | genome size | Lynch et al. (2023) |
| **B. mysticetus** | mutation rate | Lynch et al. (2023) |
| | age first reproduction | George et al. (2021b) |
| | age last reproduction | Breed et al. (2024) |
| | generation time | N/A |
| | temperature | George et al. (2021a) |
| | # of cell divisions | N/A |
| | dry mass | Lynch et al. (2023) |
| | genome size | Lynch et al. (2023) |
